# Rise-to-threshold and dynamical systems views of proactive inhibition

**DOI:** 10.1101/2021.01.16.426928

**Authors:** Vishal Rawji, Sachin Modi, Lorenzo Rocchi, Marjan Jahanshahi, John C. Rothwell

## Abstract

Successful models of movement should encompass the flexibility of the human motor system to execute movements under different contexts. One such context-dependent modulation is proactive inhibition, a type of behavioural inhibition concerned with responding with restraint. Whilst movement has classically been modelled as a rise-to-threshold process, there exists a lack of empirical evidence for this in limb movements. Alternatively, the dynamical systems view conceptualises activity during motor preparation as setting the initial state of a dynamical system, that evolves into the movement upon receipt of a trigger. We tested these models by measuring how proactive inhibition influenced movement preparation and execution in humans. We changed the orientation (PA: postero-anterior and AP: antero-posterior flowing currents) and pulse width (120 μs and 30 μs) of motor cortex transcranial magnetic stimulation to probe different corticospinal interneuron circuits. PA and AP interneuron circuits represent the dimensions of a state space upon which motor cortex activity unfolds during motor preparation and execution. AP_30_ inputs were inhibited at the go cue, regardless of proactive inhibition, whereas PA_120_ inputs scaled inversely with the probability of successful inhibition. When viewed through a rise-to-threshold model, proactive inhibition was implemented by delaying the trigger to move, suggesting that motor preparation and execution are independent. A dynamical systems perspective showed that proactive inhibition was marked by a shift in the distribution of interneuron networks (trajectories) during movement execution, despite normalisation for reaction time. Viewing data through the rise-to-threshold and dynamical systems models reveal complimentary mechanisms by which proactive inhibition is implemented.

**Key points:**

- We view proactive inhibition through the rise-to-threshold and dynamical systems models.
- We change the orientation (PA: postero-anterior and AP: antero-posterior flowing currents) and pulse width (120 μs and 30 μs) of transcranial magnetic stimulation to probe interneuron networks in motor cortex during behavioural tasks employing proactive inhibition.
- When viewed through a rise-to-threshold model, proactive inhibition was implemented by delaying the trigger to move, suggesting that motor preparation and execution are independent.
- A dynamical systems perspective showed that despite normalisation for reaction time, the trajectory/balance between PA_120_ and AP_30_ interneuron inputs during movement execution depended on proactive inhibition.
- Viewing data through the rise-to-threshold and dynamical systems models reveal complimentary mechanisms by which proactive inhibition is implemented.

## Introduction

Normal human functioning relies on movements to be made with a degree of flexibility depending on context. For example, movements can be made slower when accuracy is prioritised, captured by the speed-accuracy tradeoff (FITTS, 1954). A successful model linking neural activity to behaviour should encompass the flexibility of the human motor system to execute movements under different contexts. One such context-dependent modulation of movement is termed proactive inhibition: a prospective and goal-oriented type of behavioural inhibition concerned with responding with restraint, for example, driving slower than normal around a school in anticipation of children running out into the road (Jahanshahi *et al.*, 2015; Jahanshahi & Rothwell, 2017).

Traditionally, movement has been modelled as a rise-to-threshold process, where activity during movement preparation builds up to a threshold at a particular rate, after which movement is triggered, thereby coupling processes of motor preparation and execution. The rise-to-threshold model predicts that neural activity during movement preparation is a subthreshold form of the movement itself; although evidence for this exists in the oculomotor system (Hanes & Schall, 1996), there is a lack of empirical evidence for this hypothesis in limb movements. In fact, activity of single neurons in motor cortex (M1) during movement preparation differ greatly from activity of those same neurons during movement execution (Churchland *et al.*, 2012).

An alternative view is the dynamical systems view of motor control (Shenoy *et al.*, 2013; Vyas *et al.*, 2020). This proposes that, instead of representing explicit features of the movement (such as direction or velocity), M1 activity during motor preparation sets the initial state of a dynamical system, that evolves into the desired movement (Churchland *et al.*, 2010) upon the receipt of some trigger to move (Kaufman *et al.*, 2016). Consequently, neural activity during movement preparation and execution reflects the transition from one state to the next under some dynamical rule, and hence not all M1 activity need represent movement-related activity. Crucially, the dynamical system arises as an interplay between populations of neurons during motor preparation and execution and is not appreciated from the single-neuron perspective, which has traditionally driven theories of motor control.

The dynamical systems view uses state space models to visualise population-level neural activity. This is performed by treating each neuron’s activity as an individual axis in multi-dimensional space. A point in this space determines the state of neural population activity at a particular time. By plotting these points throughout time, a trajectory is drawn, which determines the change of neural population state throughout time (Vyas *et al.*, 2020). These states and trajectories reflect important features of movement dynamics and behaviour such as parsing motor preparation and execution into two discrete processes with independent, putative dynamics. Moreover, neural activity closer to optimal preparatory neural states results in faster reaction times (Ames *et al.*, 2014).

The investigation of dynamical systems in motor control in healthy humans has been limited by the lack of recordings from single neurons. To overcome this limitation, we leveraged a well-known property of transcranial magnetic stimulation (TMS), a non-invasive brain stimulation tool that can activate underlying cortical neurons in a focal manner. By changing the coil orientation (Mills *et al.*, 1992; D’Ostilio *et al.*, 2016; Rawji *et al.*, 2018) (PA: postero-anterior flowing current and AP: antero-posterior flowing current) and pulse width (120 μs and 30 μs) of TMS applied to M1 (D’Ostilio *et al.*, 2016; Hannah & Rothwell, 2017; Casula *et al.*, 2018; Hannah *et al.*, 2020), different corticospinal interneuron circuits can be probed. Akin to the case for single neuron activity, the activity in PA and AP interneuron circuits represent the dimensions of a state space upon which M1 activity unfolds during motor preparation and execution.

We set out to test the rise-to-threshold and dynamical systems models of movement by measuring how proactive inhibition influences movement preparation and execution in healthy humans. We do so by measuring how M1 activity, measured as corticospinal excitability (CSE), changes during movement preparation and execution during behavioural tasks that require varying degrees of proactive inhibition. Specifically, we asked whether proactive inhibition would manifest as a slower rise in CSE prior to movement onset (as predicted by rise-to-threshold models) and whether proactive inhibition would selectively affect PA or AP inputs to M1 (as predicted by dynamical systems models).

## Methods

### Participants

16 healthy volunteers (9 males, 16 right-handed) aged 19-33 (mean age 24.65, SD 4.13) participated. The study was approved by the UCL Ethics Committee and informed consent was obtained from all participants. None of the participants had contraindications to TMS, which was assessed by a TMS screening questionnaire.

### Electromyography recordings

Throughout the experiment, participants were seated comfortably in a non-reclining chair, with their right index finger rested over the ‘M’ key on a keyboard. Their forearms were supported using a cushion. Electromyographic (EMG) activity was recorded from the right first dorsal interosseous (FDI) muscle using 19 × 38 mm surface electrodes (Ambu WhiteSensor 40713) arranged in a belly-tendon montage. The raw signals were amplified, and a bandpass filter was also applied (20 Hz to 2 kHz, Digitimer, Welwyn Garden City, United Kingdom). Signals were digitised at 5 kHz (CED Power 1401; Cambridge Electronic Design, Cambridge, United Kingdom) and data were stored on a computer for offline analysis (Signal version 5.10, Cambridge Electronic Design, United Kingdom).

### Transcranial magnetic stimulation

MEPs in the right FDI muscle were evoked using a controllable TMS (cTMS) device (cTMS3, Rogue Research Inc., Canada), connected to a standard figure-of-eight coil (wing diameter 70 mm, Magstim, United Kingdom). The hotspot was identified as the area on the scalp where the largest and most stable MEPs could be obtained for the right FDI muscle, using a given suprathreshold intensity. TMS applied in postero-anterior (PA) or antero-posterior (AP) orientations allows for different interneuron inputs to a common motor (M1) output to be probed. Recent developments show that changing pulse width of TMS can allow for more selective assessment of each of the interneuron inputs probed by PA or AP TMS (Hannah & Rothwell, 2017). To this end, we delivered TMS in two ways. With the coil held approximately perpendicular to the presumed central sulcus and tangentially to the skull, TMS was given either with the coil handle pointing backwards for PA stimulation at 120 μs pulse width (PA_120_) or with the coil handle pointing forwards for AP stimulation at 30 μs pulse width (AP_30_).

### Stop-signal task and Go-only simple reaction time task

Participants were asked to perform two blocks of the stop-signal task (SST) and two blocks of a simple reaction time (Go-only) task, which were driven by custom-made MATLAB (MathWorks) scripts using Psychtoolbox. For the SST, participants were first presented with a white fixation cross on a black background. After 500 ms, an imperative stimulus (right arrow) was presented, which instructed the participant to press the ‘M’ key on the keyboard as fast as possible with their right index finger (go trials, n=105). On 25% of trials, a stop signal (red cross) appeared above the imperative stimulus at a variable delay after the imperative stimulus (stop trial, n=35). This delay, known as the stop signal delay (SSD), was controlled by a dynamic tracking algorithm, whereby the SSD would change depending on the outcome of the previous stop trial. The starting SSD was always set at 150 ms. If the participant successfully prevented their button press on a stop trial, the next stop trial would have its SSD set 50 ms later, whereas if the participant failed to stop, the next stop trial would have its SSD set 50 ms earlier. This dynamic tracking algorithm has been shown to reliably induce a convergence onto 50% successful inhibition across participants. The SSDs ranged from 100-250 ms (100, 150, 200 and 250 ms). There were also 15 baseline trials where TMS was given without cue signals to assess baseline CSE. These trials also served as catch trials. The order of trials was pseudorandomised, such that one in every four trials contained a stop signal. The Go-only reaction time task was similar to the SST, except no stop signals appeared in the block. Consequently, less proactive control was required during the Go-only task. 105 go trials were given, with 15 trials with no imperative signals to act as baseline (Figure 1). Inter-trial interval was set to 1750 ms.

**Figure 1:**
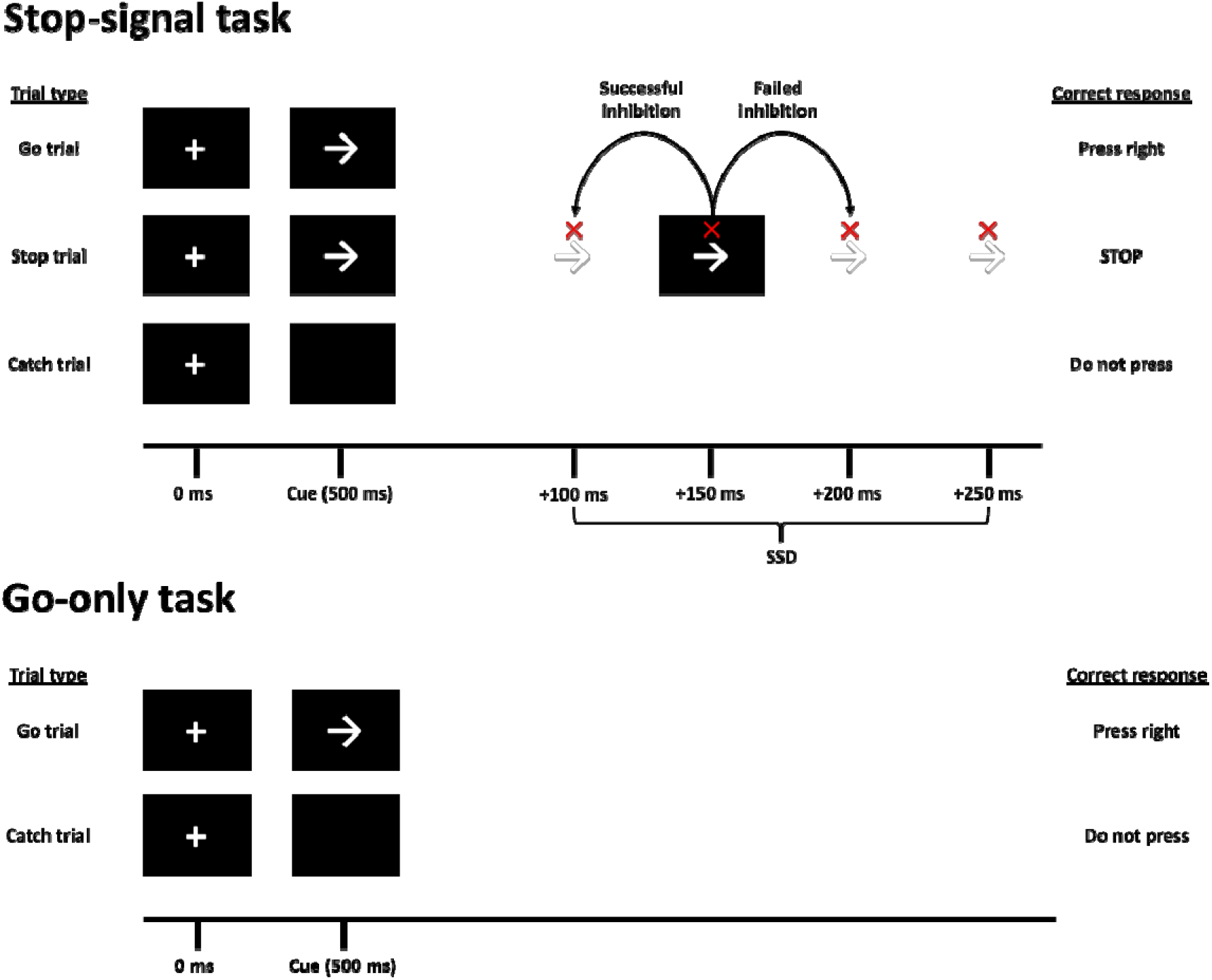
The Stop-signal and Go-only tasks.

The main behavioural measure of interest was the response delay effect (RDE) – a reaction time measure of proactive inhibition. This was calculated as the difference in reaction time on go trials in the SST and Go-only task. Other behavioural measures collected included Go reaction time (reaction time on go trials), Stop Respond reaction time (reaction time on failed stop trials), average SSD and p(inhibit) (proportion of correct stop trials on the SST). We also calculated the stop-signal reaction time (SSRT) using the mean method (Verbruggen & Logan, 2009) (mean go reaction time – mean SSD). The SSRT is a measure of reactive inhibition.

### Integration of TMS with the stop-signal and Go-only simple reaction time tasks

TMS was given on all trials, in all blocks, to the M1 representation for the right FDI muscle, at an intensity required to produce a test MEP of 0.5 mV peak-to-peak amplitude. During go trials, one TMS pulse was given randomly at one of seven time points (at the go cue and 50, 100, 150, 200, 250 and 300 ms after the go cue). 15 MEPs were taken at each time point. In the 15 baseline trials, TMS was given 1000 ms into the beginning of the trial to assess CSE at rest.

SST: Go trials consisted of presentation of a fixation cross, followed by a go cue (right arrow) 500 ms later. In 25% of trials, the right arrow was followed by a stop-signal (red cross) at one of four SSDs (100, 150, 200 or 250 ms after the arrow). Participants attempted to abort their button press on presentation of a stop-signal. Failure to do so resulted in the next stop-signal having a shorter SSD (−50 ms) whereas successful stopping led to the next SSD becoming longer (+50 ms). PA_120_ or AP_30_ TMS was delivered on go trials at one of seven time points (counterbalanced and randomised), or 1000 ms into a trial where no signals are shown (baseline trial). Go-only task: comprised of go and catch trials only; TMS was delivered at the same timepoints described above.

### Data analysis

Trials with reaction times exceeding 1000 ms were classed as omission errors due to lapses in concentration. The magnitude of proactive inhibition was determined as the reaction time difference on go trials between the SST and Go-only task, also known as the RDE. We used a paired t-test to test whether the go reaction times for the SST and the Go-only tasks were statistically different.

MEPs were pre-processed using visual inspection. Trials where TMS arrived during or after the EMG burst were excluded from analysis. To track the progression of CSE from different M1 inputs under different stopping conditions, a three-way repeated measures ANOVA with factors COIL ORIENTATION (PA_120_, AP_30_), BLOCK TYPE (SST, Go-only) and TIME POINT (Cue, 50, 100, 150, 200, 250, 300 ms) was performed using peak-to-peak MEP amplitude as the dependent variable. Post-hoc paired t-tests were then performed between MEPs at each time point against that at baseline. We also represented CSE between stopping conditions and inputs from the viewpoint of movement execution. To do this, we calculated the time between TMS delivery and response, then grouped MEPs according to 50 ms time bins from the response (300-350, 250-300, 200-250, 150-200, 100-150 and 50-100 ms before movement). Upon visually inspecting the data, we observed that the response-locked curves were remarkably similar between conditions. Given this similarity, we decided to test for equivalence between the response-locked CSE profiles using a Bayesian ANOVA. The resulting Bayes factors (BF) were used to quantify the similarity between CSE profiles (Wetzels *et al.*, 2011). Bayesian approaches are thought to be better suited to tests of equivalence than frequentist approaches (Quintana & Williams, 2018; van Ravenzwaaij *et al.*, 2019, Anon, 2020).

To further investigate the effect of proactive inhibition on behaviour and M1, we performed a detailed analysis of trial-to-trial variations in p(inhibit), reaction time and PA_120_/AP_30_ CSE. This was based on an assumption that participants would not expect successive stop signals. Therefore, we predicted that p(inhibit) and reaction time would be lowest on go trials immediately after stop trials, and that behaviour would scale as a function of the number of successive go trials. That is, participants would respond slower and p(inhibit) would increase as their expectation of a stop signal increased (with more successive go trials). We hypothesised that this may be reflected in CSE of interneuron inputs too. To this end, we used ANOVAs to quantify the relationship between successive go trials on behaviour/CSE; the main factor was GO TRIAL AFTER STOP TRIAL (one, two, three four), with the dependent variable changing as per the ANOVA (p(inhibit), reaction time). Note for CSE analyses, we used a two-way ANOVA with main factors COIL ORIENTATION (PA_120_, AP_30_) and GO TRIAL AFTER STOP TRIAL (one, two, three four) and the dependent variable being MEP amplitude. We only used MEPs from the Cue, 50 and 100 ms, given that this period is reflective of movement preparation rather than movement execution (Haith *et al.*, 2016). To further explore the relationship between behaviour and CSE, we performed Spearman’s rank correlation coefficients between p(inhibit) and reaction time, and p(inhibit) and CSE (PA_120_/AP_30_).

To examine CSE during movement preparation and execution from a dynamical systems perspective, we treated PA_120_ and AP_30_ inputs as dimensions on a single plane. That is, we plotted normalised to baseline PA_120_ MEPs (x-axis) against normalised to baseline AP_30_ MEPs (y-axis). Points on the plot show PA_120_ and AP_30_ MEPs at each time point for cue-locked (Cue, 50, 100, 150, 200, 250, 300 ms) and response-locked (300-350, 250-300, 200-250, 150-200, 100-150 and 50-100 ms before movement) analyses.

## Results

### Physiological measurements

No significant differences were found between the amplitudes of the baseline MEPs across sessions or between PA_120_ and AP_30_ conditions. As expected, AP_30_ TMS test intensities were higher than those for PA_120_ stimulation and baseline MEPs could not be elicited for three participants. Consequently, 16 participants provided data for PA_120_ TMS, 13 for AP_30_ TMS.

### Behavioural measures

Behavioural measurements in the SST and Go-only simple reaction time task are shown in Table 1. In the SST, the dynamic tracking algorithm correctly resulted in a convergence of successful inhibition in approximately 50% of trials. There was an expected significant go reaction time difference between the SST and Go-only simple reaction time blocks; a mean difference of 103.24 ms, due to the anticipation of stopping in the former (t = 7.58, p < 0.001, d = 3.07). This reaction time difference, the RDE was the behavioural manifestation of proactive inhibition.

**Table 1:**
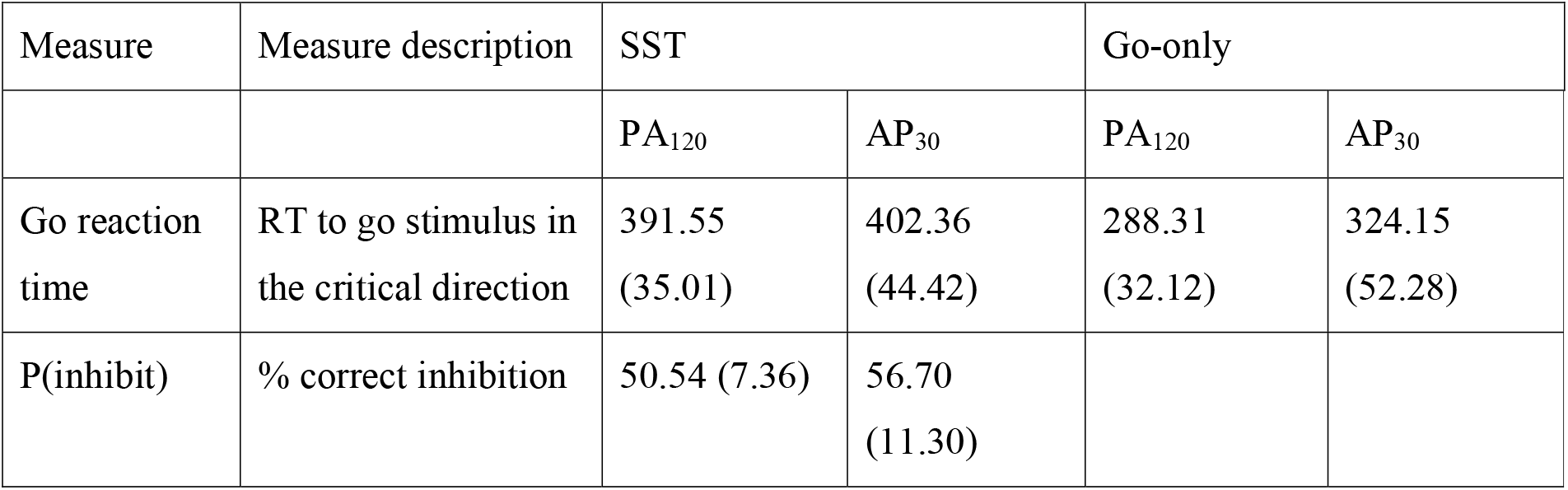
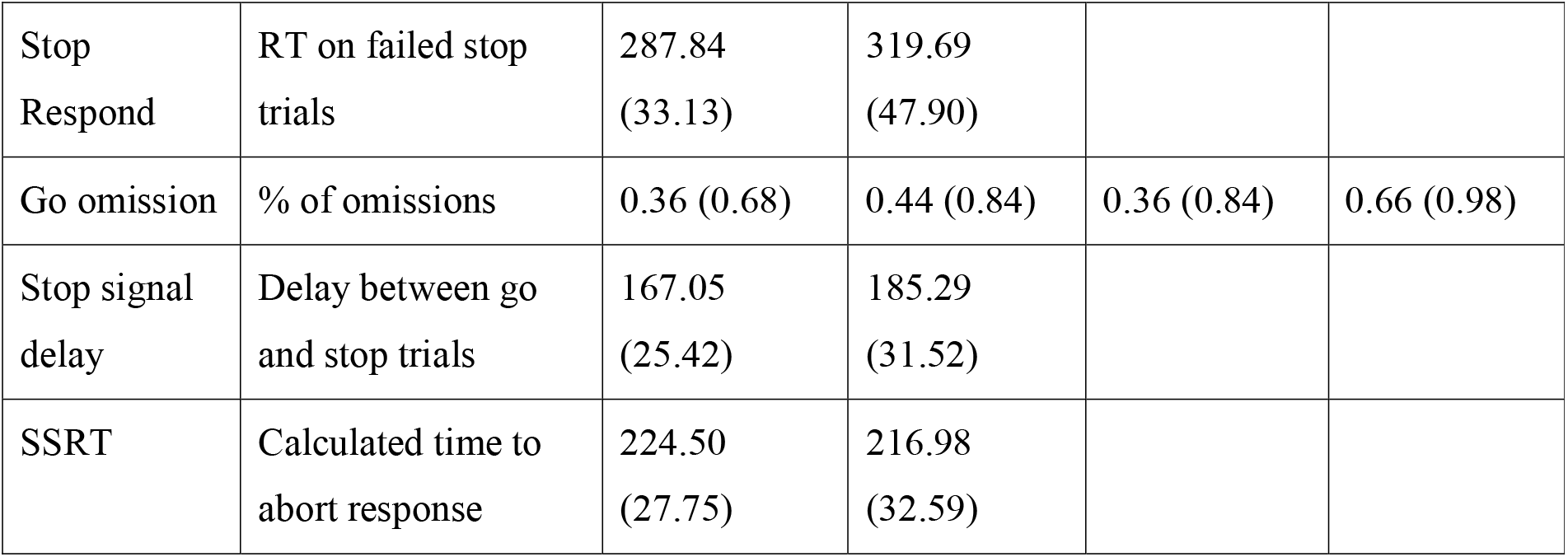
Behavioural measurements from the SST and Go-only simple reaction time tasks.

| Measure | Measure description | SST |  | Go-only |  |
| --- | --- | --- | --- | --- | --- |
|  |  | PA<sub>120</sub> | AP<sub>30</sub> | PA<sub>120</sub> | AP<sub>30</sub> |
| Go reaction time | RT to go stimulus in the critical direction | 391.55<br>(35.01) | 402.36<br>(44.42) | 288.31<br>(32.12) | 324.15<br>(52.28) |
| P(inhibit) | % correct inhibition | 50.54 (7.36) | 56.70<br>(11.30) |  |  |
| Stop Respond | RT on failed stop trials | 287.84<br>(33.13) | 319.69<br>(47.90) |  |  |
| Go omission | % of omissions | 0.36 (0.68) | 0.44 (0.84) | 0.36 (0.84) | 0.66 (0.98) |
| Stop signal delay | Delay between go and stop trials | 167.05<br>(25.42) | 185.29<br>(31.52) |  |  |
| SSRT | Calculated time to abort response | 224.50<br>(27.75) | 216.98<br>(32.59) |  |  |

The table shows the behavioural measures from the SST, Go-only simple reaction time task. Measures are accompanied by SD in brackets. Reaction times are given in ms. SSRT = stop signal reaction time

### Evolution of corticospinal excitability in stop-signal and Go-only simple reaction time tasks: cue-locked analysis

The SST was used to probe the temporal dynamics of CSE changes during which proactive inhibition is implemented (Rawji *et al.*, 2020*a*). This was compared to the same TMS timings in a task where less proactive inhibition should be employed during the Go-only simple reaction time task. We first assessed how CSE changed with respect to time by performing a stimulus-locked analysis. The three-way repeated measures ANOVA with factors COIL ORIENTATION, BLOCK TYPE and TIME POINT revealed significant effects for COIL ORIENTATION (p = 0.002, F(1,12) = 15.86, η^2^ = 0.57), BLOCK TYPE (p = 0.002, F(1,12) = 15.73, η^2^ = 0.57), TIME POINT (p < 0.001, F(7,84) = 32.51, η^2^ = 0.73) and a significant COIL ORIENTATION*BLOCK TYPE*TIME POINT interaction (p = 0.027, F(7,84) = 2.41, η^2^ = 0.17). There were no other significant effects.

### Preparation of movement

In subsequent analyses, to unravel the significant three-way interaction, data for AP_30_ and PA_120_ stimulation were treated separately. Baseline MEP sizes between go and stop blocks did not differ for PA_120_ or AP_30_ TMS, indicated by a two-way repeated measure ANOVA with factors COIL ORIENTATION and BLOCK TYPE, which revealed no significant interactions.

We compared CSE at the time of the cue with CSE at baseline. In doing so, we observed suppression of AP_30_ inputs during movement preparation in both the Go-only (Cue: p = 0.085, t = 1.46, d = 0.41; 50 ms: p = 0.046, t = 1.83, d = 0.51; 100 ms: p = 0.374, t = 0.33, d = 0.09) and SST (Cue: p = 0.027, t = 2.13, d = 0.59; 50 ms: p = 0.004, t = 3.19, d = 0.88; 100 ms: p < 0.001, t = 3.95, d = 1.10), which was not present for PA_120_ inputs (Go-only – Cue: p = 0.654, t = 0.40, d = 0.10; 50 ms: p = 0.829, t = 0.98, d = 0.25; 100 ms: p = 0.960, t = 1.88, d = 0.47; SST – Cue: p = 0.195, t = 0.88, d = 0.22; 50 ms: p = 0.050, t = 1.75, d = 0.44; 100 ms: p = 0.243, t = 0.72, d = 0.18). In addition, CSE shortly after cue presentation was lower in SST trials than in Go-only trials for PA_120_ (50 ms: p = 0.048, t = 2.16, d = 0.54; 100 ms: p = 0.008, t = 3.05, d = 0.76) and less clearly for AP_30_ stimulation (50 ms: p = 0.292, t = 1.10, d = 0.31; 100 ms: p = 0.019, t = 2.72, d = 0.75). These are summarised by plots in the top row of Figure 2.

**Figure 2:**
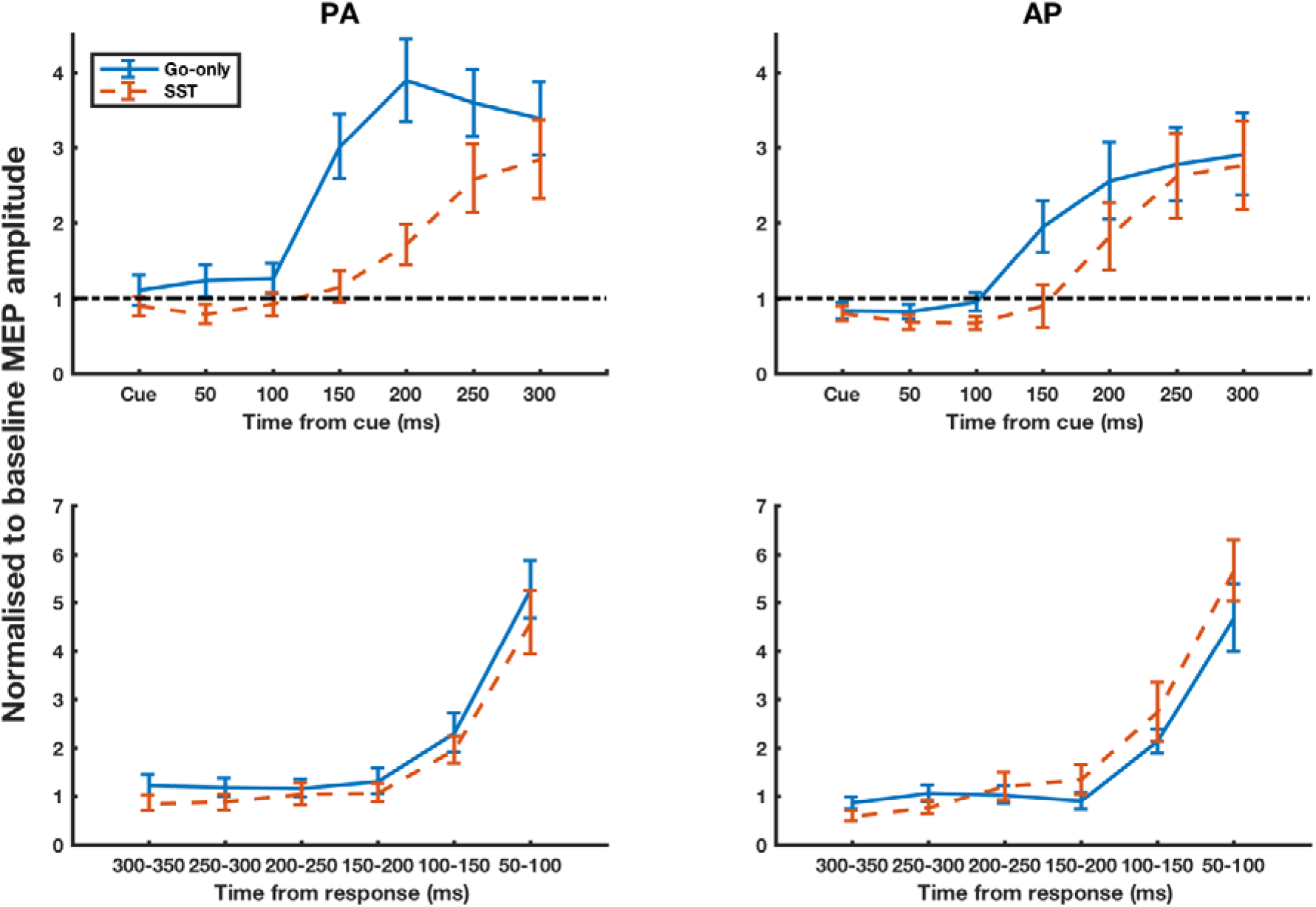
Corticospinal excitability changes during the SST and Go-only task for AP_30_ and PA_120_ TMS. Top row: MEPs are taken in go trials at various times after the go cue has been presented, for the Go-only simple reaction time task and SST. Bottom row: MEPs are sorted into 50 ms bins prior the response. MEP values are normalised to baseline MEP value. Graphs represent responses evoked using PA_120_ TMS (left column) and AP_30_ TMS (right column). Error bars represent ±SEM, Go-only task: blue solid line, SST: red dashed line.

### Execution of movement: response-locked analysis

In go trials within the SST, the main rise in excitability, indexed by the timepoint at which CSE became significantly greater than CSE at the cue, occurred later than in Go-only trials for both PA_120_ (Go-only: 100 ms, p = 0.048, t = 2.15, d = 0.39; SST: 200 ms, p = 0.002, t = 3.70, d = 0.91) and AP_30_ inputs (Go-only: 150 ms, p = 0.008, t = 3.06, d = 1.05; SST: 200 ms, p = 0.008, t = 3.04, d = 0.98). However, there was a reaction time difference between go trials in the SST and Go-only task of 103.24 ms, which may have accounted for the observed differences. Consequently, we realigned the data to the time of the response onset, thereby performing a response-locked analysis (Figure 2, bottom row). Upon visualising the data, we observed that the rate of rise in excitability preceding movement was the same during go trials in both the SST and Go-only blocks. To this end, we used Bayesian statistical methods to determine whether the rise in excitability prior to movement was equivalent between COIL ORIENTATION conditions. Indeed, the BF_10_ was 0.300 for PA_120_ stimulation and 0.385 for AP_30_ stimulation, indicating moderate evidence that CSE rise was equivalent between conditions. From this, it appears that the longer reaction times in the SST compared with the Go-only task are due to a longer pause between the presentation of the cue and rise in CSE prior to movement execution. From a rise-to-threshold perspective, there is no differential effect on PA_120_ and AP_30_ inputs to M1.

### Trial-by-trial expectation of stopping

Due to the pseudorandom design of the experiment, the probability of a stop trial occurring changed as a function of consecutive go trials. In doing so, we predicted that this would change a participant’s expectation of stopping, which would manifest behaviourally and physiologically within M1. In order to test this hypothesis, we performed a more detailed analysis of the SST data.

### Trial-by-trial behavioural analyses

Because the task was designed pseudorandomly, such that one stop trial arose in every four trials, this meant that the probability of a stop trial dynamically changed throughout the task. Consequently, the more consecutive go trials that were presented, the greater the probability that the next trial would be a stop trial. Conversely, the probability of a stop trial occurring immediately after another stop trial was lowest of all trial combinations. We compared this with behavioural data of the probability of successful inhibition on a particular stop trial, p(inhibit), based on the number of preceding go trials (Figure 3, bottom right). The ANOVA for the behavioural data showed that the number of go trials preceding a stop trial significantly modulated the probability of successfully stopping on the following trial (F(3,36) = 7.34, p = 0.001, η^2^ = 0.38). That is, the probability of successfully stopping was lowest when a stop trial occurred after 0 (STOP-STOP) or 1 (STOP-GO-STOP) go trials. The ANOVA for reaction time (Figure 3, bottom left) did not significantly change with successive go trials (F(3,36) = 1.92, p = 0.143, η^2^ = 0.14), although there was a significant positive correlation between reaction time and probability of successfully inhibiting (r_s_ = 0.37, p = 0.002).

**Figure 3:**
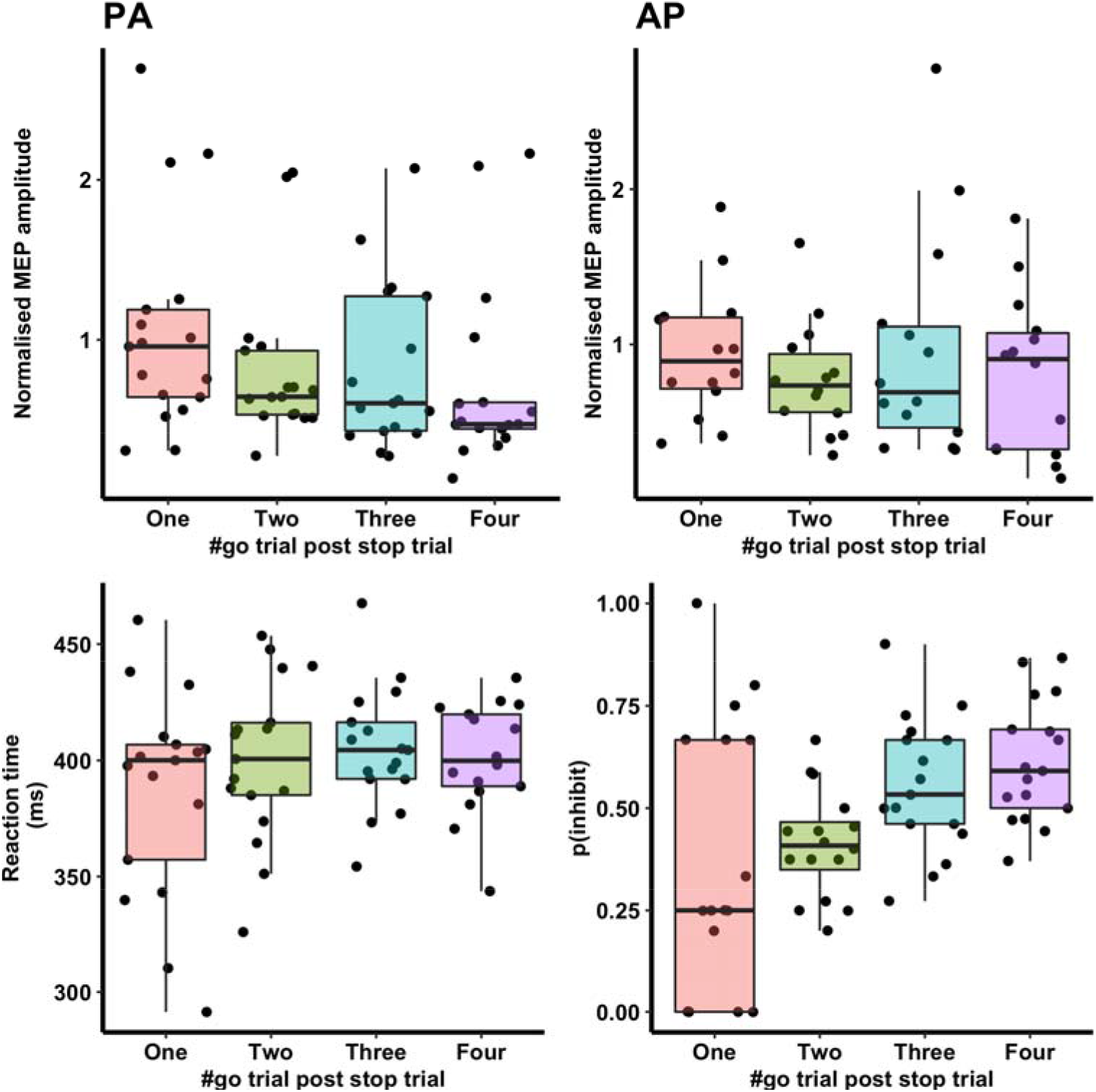
Changes in corticospinal excitability of PA_120_ and AP_30_ inputs reflected in the probability of successful inhibition and reaction times of go trials post stop trials. MEP: Top row displays mean (±SEM) normalised to baseline MEPs taken during go trials in the SST using PA_120_ and AP_30_ TMS. A grand average for each participant was measured by averaging MEPs at the cue, 50 ms and 100 ms. This is plotted against successive go trials after stop trials. Behaviour: Bottom row shows the probability of successfully inhibiting a response and reaction time on progressive go trials after a stop trial has been shown. Boxplots are coloured such that MEP measures and behaviours correspond if they are the same colour.

### Trial-by-trial corticospinal excitability analyses

We first assessed if CSE was influenced by coil orientation or number of go trials after a stop trial using a two-way repeated measures ANOVA. This showed no significant effects of COIL ORIENTATION (p = 0.894, F(1,12) = 0.02, η^2^ < 0.01), GO TRIAL AFTER STOP TRIAL (p = 0.357, F(4,48) = 1.12, η^2^ = 0.03) or an interaction between the two (p = 0.317, F(4,48) = 1.21, η^2^ = 0.02). Given that there was an unequal number of participants in each group (PA_120_ = 16 and AP_30_ = 13) and our prior hypothesis that proactive inhibition would differentially modulate interneuron inputs, we decided to perform an exploratory analysis of PA_120_ and AP_30_ inputs separately. PA_120_ MEPs were modulated depending on which go trial they were evoked on after a stop signal; MEP size was greatest for trials immediately following a stop trial and decreased with increasing number of go trials (p = 0.045, F(3,36) = 3.64, η^2^ = 0.19). Conversely, there was no such relationship present for AP_30_ MEPs (p = 0.399, F(3,36) = 1.10, η^2^ = 0.07), shown in Figure 3. To assess whether this was a true suppression in the light of potential stopping, we compared the raw CSE from PA_120_ MEPs with those collected at equivalent times during a Go-only task, where no stop signals were presented; consequently, these MEPs reflect early time point CSE when no proactive control is expressed. Paired t-tests showed significant suppression of PA_120_ MEPs, which were taken on the 2^nd^ (p = 0.031, t = −2.38, d = 0.63), 3^rd^ (p = 0.046, t = −2.18, d = 0.60) and 4^th^ (p = 0.019, t = −2.62, d = 0.73) go trials after a stop trial. Finally, whilst PA_120_ and AP_30_ MEP amplitudes were both positively correlated with reaction time (PA_120_: r_s_ = 0.27, p = 0.028; AP_30_: r_s_ = 0.32, p = 0.017), only PA_120_ MEPs were significantly correlated with the probability of successful inhibition (PA_120_: r_s_ = 0.24, p = 0.049; AP_30_: r_s_ = 0.02, p = 0.915; comparison between AP_30_ and PA_120_ correlations, p = 0.06).

### Proactive inhibition from a dynamical systems perspective

We plotted normalised to baseline PA_120_ and AP_30_ MEPs against each other for each timepoint in the cue and response-locked analyses (Figure 4). The resultant trajectories show how population-level activity within M1 evolves during movement preparation and execution. The cue-locked analysis shows that M1 population activity evolves within the same subspace early after cue presentation (bottom left). Approximately 150 ms later, activity increases in both interneuron networks and occupies a separate space at the end of movement (top right). Notably, the trajectories taken vary between the tasks, although this may be due to differences in reaction times. However, the response-locked analyses similarly show a difference in trajectories, despite normalisation for reaction time. From the dynamical systems view, behaviourally equivalent responses are executed differently by M1 population activity, dependent on the requirements for proactive inhibition.

**Figure 4:**
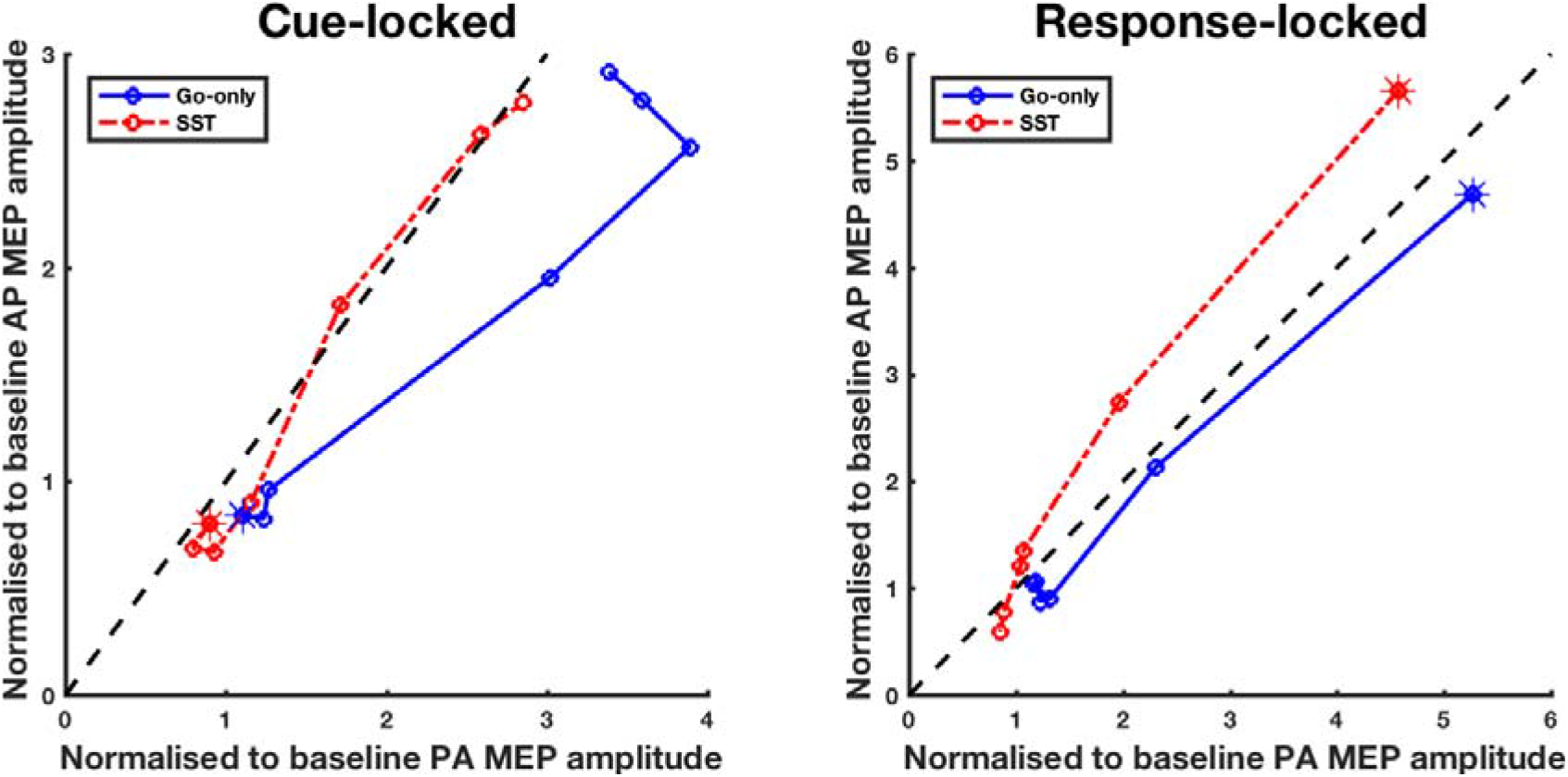
Motor cortex population-level activity during movement preparation and execution. Motor cortex population-level activity is represented as a combination between PA_120_ and AP_30_ inputs. Each plot shows the trajectory taken by this population activity throughout movement. Cue-locked analysis: activity starts at the cue (shown by stars). Activity then progresses over time, with each marker (circle) representing a time-point (Cue, 50, 100, 150, 200, 250 and 300 ms). This is shown for the SST (red, dashed line) and Go-only task (blue, solid line). Response-locked analysis: stars represent population activity 50-100 ms prior to movement. Working backwards, time-points are as shown in the bottom row for Figure 2. Dashed, x=y lines represent balanced PA_120_ and AP_30_ CSE.

## Discussion

### A rise-to-threshold view of proactive inhibition

We sought to investigate how proactive inhibition was implemented by the motor system. We used two tasks which differed in their instructions, such that participants employed greater proactive inhibition in the SST, as indexed by a prolonged reaction time (RDE) (Obeso *et al.*, 2014; Rawji *et al.*, 2020*a*, 2020*b*). By applying TMS, we observed that, relative to the time of onset of the go cue, CSE increased more slowly during the go trials of the SST compared with the Go-only task. In contrast, from the viewpoint of motor execution (response-locked analysis), the rate of increase in CSE was the same in both tasks. Hence, the delayed rise in CSE probably arose from a delay in the trigger to initiate the rise in CSE, rather than a slower rise of CSE to a notional “threshold” for movement (Rawji *et al.*, 2020*b*).

Our results reveal a dissociative role of M1 inputs during movement: whereas AP_30_ inputs are suppressed regardless of proactive inhibition (due to preparatory inhibition), PA_120_ inputs reflect some element of the expectation to go or stop as expected from proactive inhibition. In this task, the effector being called into action is always the right index finger. Therefore, we assume that motor preparation, from an effector selection perspective, is equal in the Go-only and SST. AP_30_ inputs are suppressed with respect to baseline in both tasks (Figure 2, top right) and this suppression does not change as a function of the expectation to stop (Figure 3, top right). Previous authors have noted that CSE is depressed at around the expected time of onset of the “go” cue even in simple reaction time tasks, and our studies have confirmed that this is preferentially due to effects on elements recruited by AP_30_ inputs to M1 (Hannah *et al.*, 2017; Ibáñez *et al.*, 2019). This preparatory inhibition may reflect a form of volitional proactive inhibition that prevents premature release of the intended movement; alternatively, it may represent a necessary part of preparing to move. Given that the suppression occurs regardless of proactive inhibition (RDE), it is more likely that CSE suppression at the time of the go cue represents a necessary part of movement preparation (Vassiliadis *et al.*, 2020). Hence, what is being assayed in these early time points, in AP_30_ MEPs, is probably an epiphenomenon of movement preparation – preparatory inhibition. PA_120_ inputs, on the other hand, scale with stopping probability; they are suppressed when stopping might be required, in a dose-dependent fashion. Overall, these results point to two simultaneous processes occurring: AP_30_ suppression reflecting preparatory inhibition (i.e., preparation for the forthcoming motor trigger), which is overlaid by suppression of PA_120_ inputs, reflecting the possibility of stopping – proactive inhibition.

### A dynamical systems view of proactive inhibition

An alternative explanation for the behaviour of these inputs during proactive inhibition can be understood from the dynamical systems view of motor control (Shenoy *et al.*, 2013; Vyas *et al.*, 2020). The dynamical systems approach is appropriate to interpret TMS results, given that they both rely on population-level activity. By plotting PA_120_ CSE against AP_30_ CSE, we visualise how M1 *population* activity evolves throughout time and through movement preparation and execution (Figure 4). Akin to the findings using a linear dynamical systems approach, we see that activity during movement preparation evolves in a particular, confined subspace for approximately 150 ms after cue presentation. That is, activity occupies the bottom left of the cue-locked plot for 150 ms (similarly, activity occupies the bottom left of the response-locked plot 300-350 ms prior to movement). Following this, M1 population CSE suddenly and dramatically increases, upon receipt of a trigger for movement execution, to a different area in the subspace (top right of the cue-locked and response-locked plots).

A key feature is that, during motor preparation, population activity evolves along a “null space” where the activity of individual neurons on corticospinal outputs sums to zero through varying degrees of excitation and inhibition; this results in a new state primed for motor execution but in which there is no change in corticospinal output (Kaufman *et al.*, 2014; Stavisky *et al.*, 2017). In essence, the decrement in AP_30_ inputs during movement preparation may be an epiphenomenon of the aforementioned excitation-inhibition balance that causes neural activity to evolve in the null space. In contrast, PA_120_ inputs may have a different functional role; given that they scale with the probability of successfully inhibiting and not reaction time, PA_120_ inputs may represent a “stop” process or activity in a “proactive inhibition space”. This idea is consistent with prior hypotheses regarding CSE suppression during movement preparation, which serves to reduce noise (Greenhouse *et al.*, 2015; Lebon *et al.*, 2019) so that other inputs more easily select task-relevant effectors. Although the cited studies use PA-oriented stimulation, they do so without changing pulse width and hence are less selective for PA_120_ inputs studied in the current study.

An important observation is that the trajectory taken (or balance between PA_120_ and AP_30_ inputs) during movement differs between tasks in the response-locked analysis, despite being normalised for reaction time and movement dynamics (right button press). As aforementioned, movement preparation is similar in the two tasks, as indexed by activity occupying the same subspace early in the cue-locked analysis and long before response in the response-locked analysis. The linear dynamical systems theory of movement posits that preparatory activity evolves into the movement upon receipt of a trigger to move.

Mathematically, the specific form of this trigger represents a transformation of preparatory activity into movement activity. Given that population-level preparatory activity and reaction time are equivalent between tasks but have different CSE trajectories (Figure 4, response-locked), we infer that the nature of the trigger to move (or mathematical transformation from preparatory activity to movement activity) must be sensitive to how the movement will be executed – in this case, by proactive inhibition or action restraint. The ability to shift the distribution of CSE between interneuron networks may allow for qualitatively equivalent movements to be performed in a variety of ways depending on task-specific goals. To our knowledge, this is the first visualisation of CSE during movement preparation and execution represented as the interplay between different interneuron inputs in humans.

### Independent, concurrent processes during movement in motor cortex

Another prediction from the dynamical systems view of motor control is that the preparation and execution of movements are independent processes; activity accumulated during preparation evolves into the movement upon the receipt of a trigger (Churchland *et al.*, 2012). This trigger for movement is condition-invariant such that it conveys no information about *what* the movement will be but only *when* it occurs (Kaufman *et al.*, 2016). Whilst the reaction time and cue-locked CSE differ between the SST and Go-only task, the response-locked CSE profiles are remarkably similar (Figure 2, bottom row), suggesting that movement preparation and execution are independent (Haith *et al.*, 2016). The similarity between response-locked CSE profiles between conditions also mirrors the condition-invariant signal found in population-level neural recordings (Kaufman *et al.*, 2016). Interestingly, the magnitude of this execution-related component was found to be the largest of any tuned components during movement preparation – a feature comparable to the major rise in CSE prior to movement execution in our study.

### Limitations

The pseudorandom design of the task meant that participants could develop expectancy and learn to anticipate the stop signal, which could potentially confound measures of response inhibition. However, we observed that participants successfully inhibited their responses on approximately 50% of stop trials, showing that participants correctly engaged with task demands. In light of the equivalence shown by the response-locked analyses, we concluded that participants were delaying their trigger to move. Variants of the SST have shown that proactive inhibition can sometimes be mediated by alterations in the threshold before which movement is triggered (Rawji *et al.*, 2020*a*, 2020*b*). Given these apparent differences in the strategy used to mediate proactive inhibition, it may be the case that our findings are a feature of task design and differ when proactive inhibition is mediated differently. The movement (right finger button press) was known throughout the experiment and did not change, meaning that the same movement was prepared in all conditions, in all trials. Consequently, the similarity in response-locked CSE profiles (Figure 2) may be so because the movement to be prepared is the same in both conditions (although this would not account for differences in the population-activity analysis in Figure 4. Future studies should aim to change the way in which the same movement is prepared to help establish whether movement execution CSE is dependent on motor preparation.

## Conclusions

By using subtle TMS manipulations, we show that proactive inhibition mediates its effects by altering the inputs to M1. Despite being differentially modulated during motor preparation and inhibition, we do not believe that PA_120_ and AP_30_ inputs are the exclusive pathways mediating these processes. Our interpretation is more conservative, that motor preparation and inhibition can act via different inputs and that our data strengthen the hypothesis that PA_120_ and AP_30_ inputs to M1 are physiologically and behaviourally distinct (Ni *et al.*, 2011; Hamada *et al.*, 2014; Volz *et al.*, 2015; Hannah *et al.*, 2017). Rather than being contradictory, we show that viewing data through the rise-to-threshold and dynamical systems models reveal complimentary mechanisms by which proactive inhibition is implemented: proactive inhibition is implemented by delaying the trigger to move (rise-to-threshold) and occurs through a shift in the distribution of interneuron networks (trajectories) during movement execution (dynamical systems). Future work should aim to address if the dissociated role of interneuron inputs generalises when proactive control is mediated using a different strategy.

## Acknowledgements

This study was funded by doctoral training grant MR/K501268/1 from the MRC. VR would like to thank Miss Catherine Storm for her support during the writing of this manuscript.

## Author contributions

All authors contributed to the design of the study and were involved in the drafting and revisions of the manuscript. VR and SM collected the data. All authors have approved this manuscript.

## Data availability

The data that support the findings in this study are available from the corresponding author.

